# Recovery of Consciousness and Cognition after General Anesthesia in Humans

**DOI:** 10.1101/2020.05.28.121269

**Authors:** George A. Mashour, Ben J.A. Palanca, Mathias Basner, Duan Li, Wei Wang, Stefanie Blain-Moraes, Nan Lin, Kaitlyn Maier, Maxwell Muench, Vijay Tarnal, Giancarlo Vanini, E. Andrew Ochroch, Rosemary Hogg, Marlon Schwarz, Hannah Maybrier, Randall Hardie, Ellen Janke, Goodarz Golmirzaie, Paul Picton, Andrew McKinstry-Wu, Michael S. Avidan, Max B. Kelz

## Abstract

Understanding how consciousness and cognitive function return after a major perturbation is important clinically and neurobiologically. To address this question, we conducted a three-center study of 30 healthy humans receiving general anesthesia at clinically relevant doses for three hours. We administered a pre- and post-anesthetic battery of neurocognitive tests, recorded continuous electroencephalography to assess cortical dynamics, and monitored sleep-wake activity before and following anesthetic exposure. We hypothesized that cognitive reconstitution would be a process that evolved over time in the following sequence: attention, complex scanning and tracking, working memory, and executive function. Contrary to our hypothesis, executive function returned first and electroencephalographic analyses revealed that frontal cortical dynamics recovered faster than posterior cortical dynamics. Furthermore, actigraphy indicated normal sleep-wake patterns in the post-anesthetic period. These recovery patterns of higher cognitive function and arousal states suggest that the healthy human brain is resilient to the effects of deep general anesthesia.

## Introduction

The recovery of neurocognitive function after brain network perturbations such as sleep, general anesthesia, or disorders of consciousness is of both scientific and clinical importance.

Scientifically, characterizing recovery processes after such perturbations might provide insight into the more general mechanisms by which consciousness and cognition are reconstituted after major network disruptions. The ability to recover cognitive function quickly after sleep, for example, likely confers a natural selection advantage. Moreover, understanding which brain functions are most resilient could inform evolutionary neurobiology (Mashour and Alkire, 2013; Kelz and Mashour, 2019). Clinically, understanding the specific recovery patterns after pathologic states of unconsciousness could inform prognosis or therapeutic strategies. However, it is challenging to characterize differential cognitive recovery after sleep because of the rapidity of the process, whereas it can be impossible in pathologic states because of the unpredictable recovery. General anesthesia, by contrast, represents one controlled and reproducible method by which to perturb consciousness and cognition, followed by systematic observations of the recovery process. Studying recovery of cognition after general anesthesia in humans is also of particular importance in humans because animal studies have suggested that general anesthetics have the potential to immediately and persistently impair cognition in the post-anesthetic period (Culley et al., 2004; Valentim et al., 2008; Carr et al., 2011; Callaway et al., 2012; Zurek et al., 2012; Jevtovic-Todorovic et al., 2013; Zurek et al., 2014; Avidan and Evers, 2016; Jiang et al., 2017), creating a potential public health concern for the hundreds of millions of surgical patients undergoing general anesthesia each year (Weiser et al., 2015).

To improve scientific understanding of recovery of consciousness and cognition after anesthetic-induced unconsciousness, we studied 30 healthy volunteers at three centers who were administered deep general anesthesia using isoflurane for three hours, with cognitive testing conducted at pre-anesthetic baseline as well as every 30 min for three hours after return of consciousness (**Figure 1**). We hypothesized that the pattern of post-anesthetic recovery would proceed with a return of responsiveness, followed by: attention, complex scanning and visual tracking, working memory, and lastly executive function (as assessed by, respectively, Psychomotor Vigilance Test (PVT), Motor Praxis (MP), Digit Symbol Substitution Test (DSST), fractal 2-Back (NBCK), Visual Object Learning Test (VOLT), and Abstract Matching (AM)). Isoflurane anesthesia was chosen because of its heterogeneous molecular targets, which affect multiple neural systems, and because its slower offset compared to other anesthetics would allow us to observe differential recovery of function (Hemmings et al., 2019). The three-hour duration of anesthesia was chosen based on clinical data related to recovery of surgical patients, the pharmacokinetics of isoflurane, and practical considerations for volunteers participating in day-long experiments. To control for the learning effects of repeated cognitive testing (Basner et al., 2018), we also recruited 30 healthy volunteers who, instead of receiving anesthesia, were engaged in wakeful behaviour for three hours and then underwent equivalent cognitive testing at time points corresponding to the cohort that underwent general anesthesia (**Figure 1**). All participants received actigraphy watches to monitor sleep-wake activity before and after anesthesia or the control condition, and all participants had electroencephalographic recording throughout the experiment to assess local and global cortical dynamics. With the control group serving as a reference, the aims of the study were: (1) to assess the sequence of cognitive recovery following emergence from a prolonged state of unconsciousness with serial neurobehavioral assessments to test the hypothesis that higher executive functions reconstitute only after more primitive functions; and (2) to measure correlated changes in the dynamics of functional brain networks that might account for the differential recovery of cognitive function.

**Figure 1:**
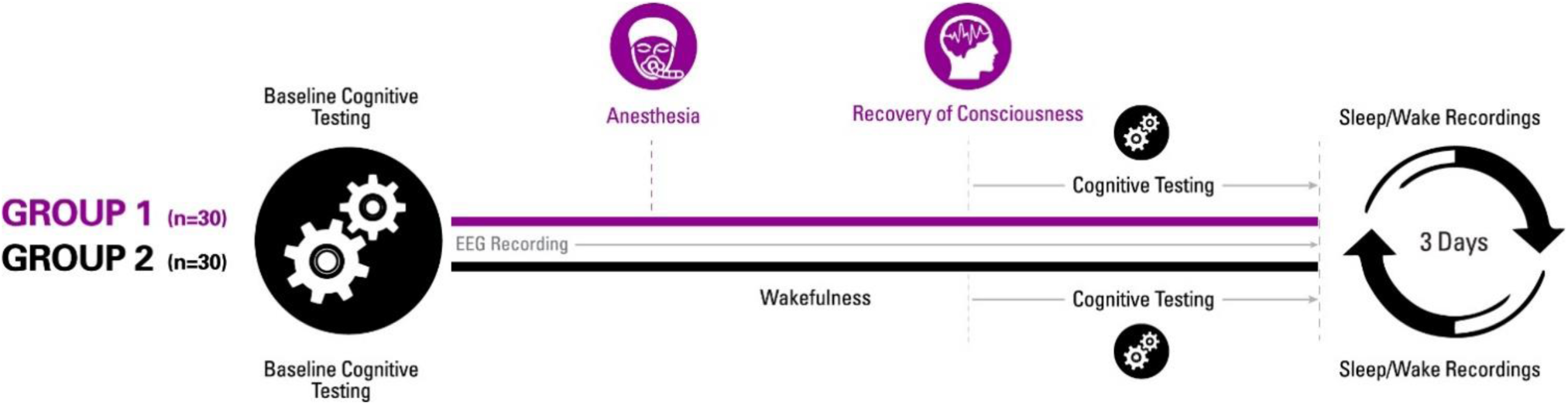
Experimental design. Participants were randomised to one of two groups for investigating recovery of consciousness and cognition after general anesthesia. Sleep-wake actigraphy data were acquired in the week leading up to the day of the experiment, which started with baseline cognitive testing followed by either a period of general anesthesia (1.3 age-adjusted minimum alveolar concentration of isoflurane) or wakefulness. Upon recovery of consciousness (or similar time point for controls), recurrent cognitive testing was performed for three hours. Actigraphy resumed for three days after the experiment.

## Results

### Participants

The study received ethics committee approval at all three sites independently; written informed consent was obtained after careful discussion with each participant. The average age of all study participants was 27 (± 4.5) years, with 50% females. There were no adverse events during the course of the study or at one-week follow up at the completion of the study.

### Recovery of cognitive functions after general anesthesia

We administered six distinct cognitive tests at baseline and twice per hour for three hours after exposure to general anesthesia or a comparable period without anesthetic exposure in the control participants. The order of the MP, PVT, DSST, NBCK, VOLT, and AM tests was randomised between subjects but consistent within subjects. In anesthetized volunteers (with correction for learning based on results from tests taken at corresponding times by the non-anesthetized controls), the speed and accuracy of all six cognitive tests were significantly impaired compared with the pre-anesthetic baseline assessments (all twelve statistical tests yielded p<0.008, with adjustments for multiple comparisons). Thus, the first question answered is that all tests were impaired at initial recovery.

The next question to be answered was whether rates of recovery differed among cognitive domains and whether the time to recovery exceeded 30 minutes when comparing among cognitive tests. Based on likelihood ratio tests, there were statistically significant differences in the rates of recovery of the six cognitive domains, both with regard to their accuracy and speed (p<0.05). Results from Bayesian analyses yielded posterior probability estimates that differences in recovery times between the various cognitive domains for accuracy and speed exceeded 30 minutes. Hence, recovery of cognition is a process that appears to evolve over time rather than one that occurs simultaneously.

For accuracy (**Figure 2**), the probability was high that (i) recovery of AM occurred more than 30 min before NBCK (95%), DSST (77%), VOLT (72%) and MP (65%); (ii) recovery of PVT occurred more than 30 min before NBCK (99.9%), DSST (81%), MP (76%) and VOLT (72%); (iii) recovery of VOLT occurred more than 30 min before NBCK (95%), DSST (76%), and MP (68%); and that (iv) recovery of MP occurred more than min before NBCK (57%).

**Figure 2:**
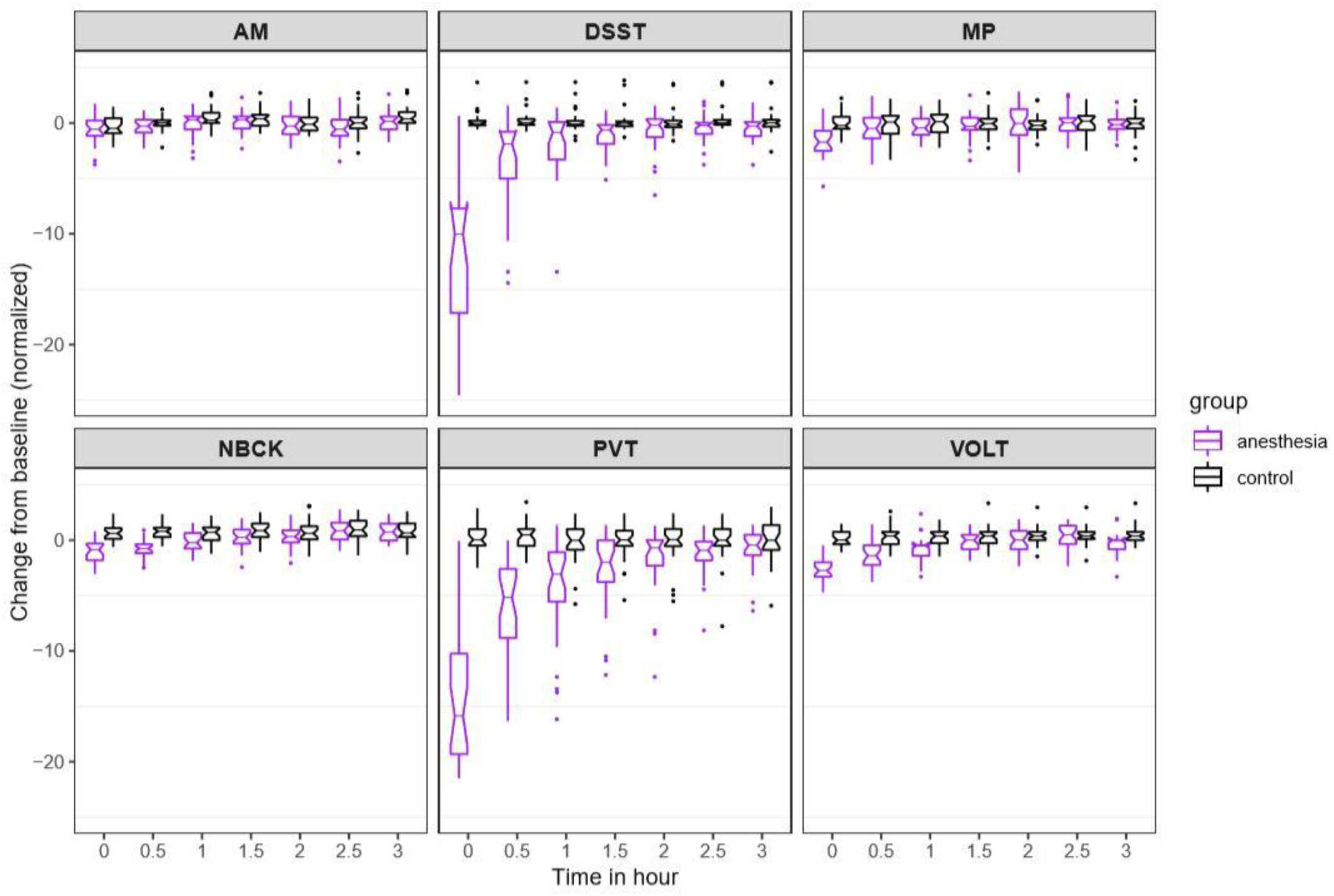
Time course for recovery of (normalized) accuracy in cognitive task performance after general anesthesia (time 0 is just after recovery of consciousness in the group that was anesthetized). AM, Abstract Matching; DSST, Digit Symbol Substitution Test; MP, Motor Praxis; NBCK, Fractal 2-Back; PVT, Psychomotor Vigilance Test; VOLT, Visual Object Learning Test. The six cognitive tests are all represented.

For speed (**Figure 3**), the probability was high that (i) recovery of AM occurred more than 30 min before PVT (68%), VOLT (64%) and DSST (52%); (ii) recovery of NBCK occurred more than 30 min before PVT (74%), VOLT (66%) and DSST (63%); (iii) recovery of MP occurred more than 30 min before PVT (81%); and that (iv) recovery of DSST occurred more than 30 min before PVT (53%). We had further hypothesized that, if there were differential rates of recovery in cognitive domains, higher order executive function (as tested by AM) would be most impaired and consequently would be the last to recover. Counter to this expectation, AM was one of the least impaired of the tests at ROC. For both speed and accuracy measures, performance on AM quickly approached its pre-anesthetic level.

**Figure 3:**
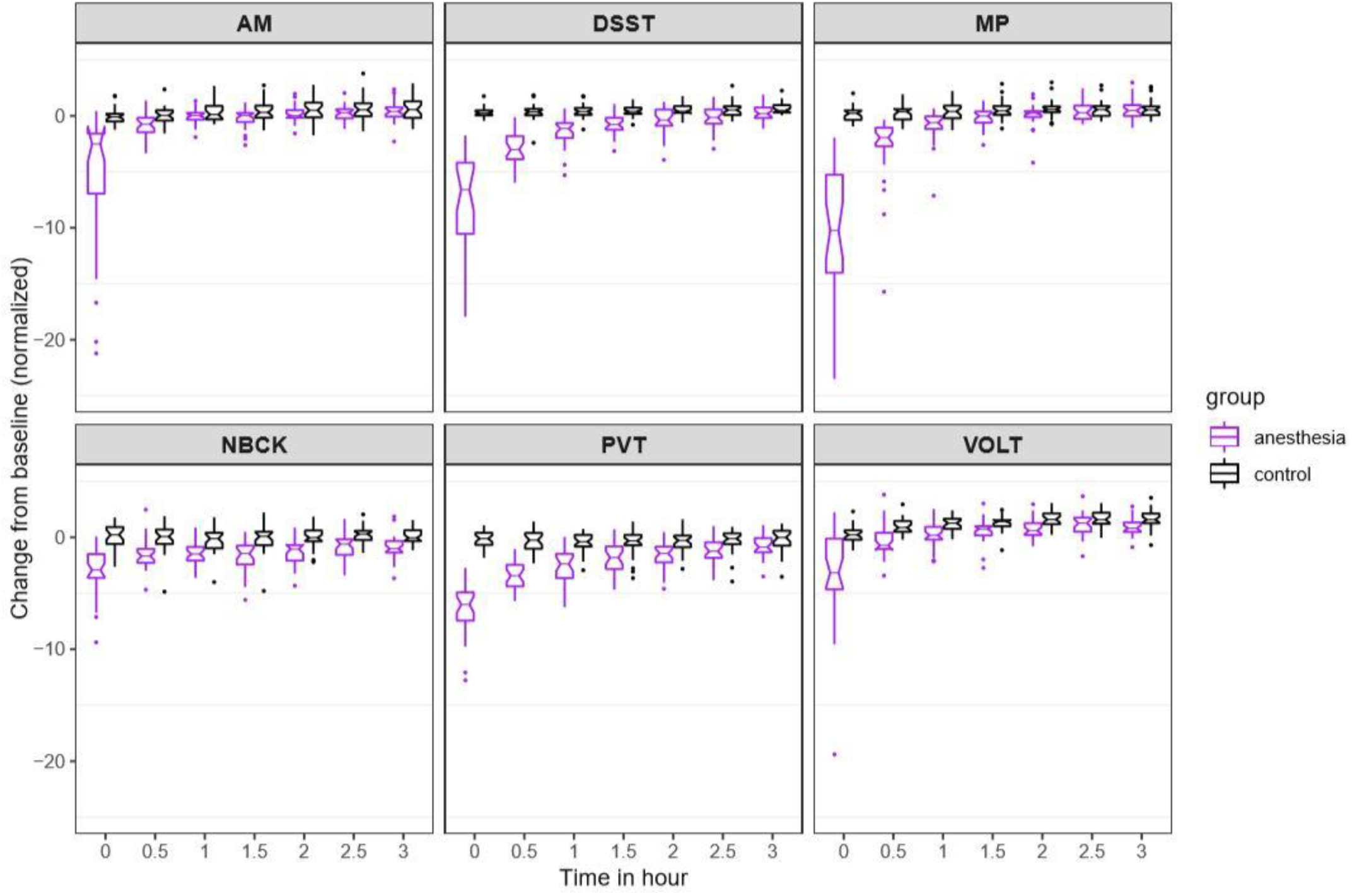
Time course for recovery of (normalized) speed of cognitive task performance after general anesthesia (time 0 is just after recovery of consciousness in the group that was anesthetized). AM, Abstract Matching; DSST, Digit Symbol Substitution Test; MP, Motor Praxis; NBCK, Fractal 2-Back; PVT, Psychomotor Vigilance Test; VOLT, Visual Object Learning Test. The six cognitive tests are all represented.

As expected, for all tests, the maximal degree of impairment was upon ROC. Information on the trajectory of recovery for an individual cognitive test is depicted in Figures 2 and 3, and was assessed using a non-linear mixed effects model. In the 3-hour follow-up period, accuracy for those in the anesthesia group increased gradually for some tests (DSST, PVT) and more rapidly for others (AM, NBCK, MP, VOLT). At three hours after ROC in the anesthetized group, accuracies in five out of six tests were not statistically significantly different from the control group. There remained a significant (p<0.001), albeit small, difference in accuracy performance between the anesthetized and the control group on the VOLT at three hours. Overall, within three hours of return of consciousness, the anesthetized group returned to an accuracy level that was not substantially different from that of participants who were not anesthetized. The comparison with the non-anesthetized control group is highly informative, because assessing performance on cognitive tests over time is confounded by learning. Had the anesthetized group simply returned to, or even exceeded, their own baseline performance after three hours of testing, this would not have provided sufficient evidence to conclude that their cognition had truly returned to baseline. The inclusion of an awake control group, however, strengthens the conclusion that the anesthetized group did not experience a decrement in their performance that might have been masked by learning, which is known to occur with repeated testing.

To account for a trade-off in accuracy versus speed, we also evaluated speed of task performance (**Figure 3**). Speed was also most strongly impaired at ROC, with a drop of more than 5 SD for all tests but the NBCK. At three hours after ROC in the anesthetized group, speed in two (MP and PVT) out of six tests were not statistically significantly different from the control group. There remained significant (p<0.001), albeit small, differences between the anesthetized and the control group in speed performance on four (VOLT, NBCK, AM, DSST) tests at three hours. The results of the nonlinear mixed-effects models for cognitive performance in anesthetized and non-anesthetized cohorts are summarized in Table 1.

**Table 1:** Results of nonlinear mixed-effects models comparing cognitive trajectories at three hours post emergence between anesthetized and non-anesthetized cohorts. AM, Abstract Matching; DSST, Digit Symbol Substitution Test; MP, Motor Praxis; NBCK, Fractal 2-Back; PVT, Psychomotor Vigilance Test; VOLT, Visual Object Learning Test. For speed and accuracy of each task, we report the Bonferroni corrected confidence intervals (CI) of the difference with an overall significance level 0.05. The P-values of testing difference between groups should be compared to the corrected level 0.05/6=0.0083.

|  | SPEED |  |  | ACCURACY |  |  |
| --- | --- | --- | --- | --- | --- | --- |
|  | Estimate | P-value | CI | Estimate | P-value | CI |
| MP | 0.3562 | 0.1101 | ( -0.2437, 0.9561 ) | 0.0366 | 0.8721 | ( -0.5824, 0.6556 ) |
| VOLT | 0.7666 | 0.0720 | ( -0.3757, 1.9089 ) | 0.3915 | 0.0847 | ( -0.2184, 1.0015 ) |
| NBCK | 1.2803 | <0.0001 | ( 0.6642, 1.8964 ) | 0.0089 | 0.9664 | ( -0.5679, 0.5857 ) |
| AM | 0.6336 | 0.0023 | ( 0.09021, 1.1770 ) | 0.3692 | 0.0709 | ( -0.1791, 0.9176 ) |
| PVT | 0.7986 | 0.0026 | ( 0.1052, 1.4921 ) | 1.0916 | 0.0557 | ( -0.4359, 2.6191 ) |
| DSST | 1.0184 | <0.0001 | ( 0.4420, 1.5949 ) | 0.5454 | 0.0931 | ( -0.3279, 1.4188 ) |

### Cortical dynamics before, during, and after general anesthesia

We assessed cortical dynamics before, during, and after anesthetic exposure using local measures of permutation entropy (PE) and global measures of Lempel-Ziv complexity (LZC). The PE demonstrated significant differences associated with behavioural states (F_9, 86_ = 42.423, p<0.001), brain regions (F_1, 257_ = 4.275, p=0.040) and the interaction between them (F_9, 85_ = 2.750, p=0.007). As compared to the baseline condition of eyes-closed resting state, frontal PE decreased at propofol-induced loss of consciousness (LOC), further decreased during maintenance of the anesthetized state with isoflurane anesthesia (p<0.001, −0.160 (−0.185 to - 0.135), maintenance vs. EC1), and returned to or even exceeded the baseline level just before the recovery of consciousness (ROC) (p=0.002, 0.036 (0.018 to 0.055), pre-ROC vs. EC1) (**Figure 4A and B**). Posterior PE did not show significant changes at LOC but was decreased during the maintenance phase (p<0.001, −0.110 (−0.135 to −0.085), maintenance vs. EC1) and then returned to baseline level just before ROC (**Figure 4A and B**). The topographic maps of PE exhibited region-specific patterns, in which frontal channels demonstrated significantly higher PE values as compared to posterior channels at the eyes-closed resting state directly after emergence (EC2, p=0.002, 0.032 (0.016 to 0.048), frontal vs. posterior) (**Figure 4A**). The LZC demonstrated significant state-dependent differences (F_9, 50_ = 59.364, p<0.001). As compared to baseline consciousness, LZC declined at LOC and decreased further during the maintenance phase (p<0.001, −0.258 (−0.285 to −0.231), maintenance vs. EC1), returning to baseline level just before ROC (**Figure 4C**). Thus, both the local (PE) and global (LZC) metrics of cortical dynamics recover just as the brain is recovering consciousness. Similar state-dependent changes were observed despite different strategies in the parameter selection for PE (**Figure S1**), the threshold of binarization, and the method of generating surrogate data for LZC (**Figure S2**). Additionally, as expected, the non-anesthetized control group showed no differences among the seven resting-state eyes-closed epochs (**Figure S3**).

**Figure 4:**
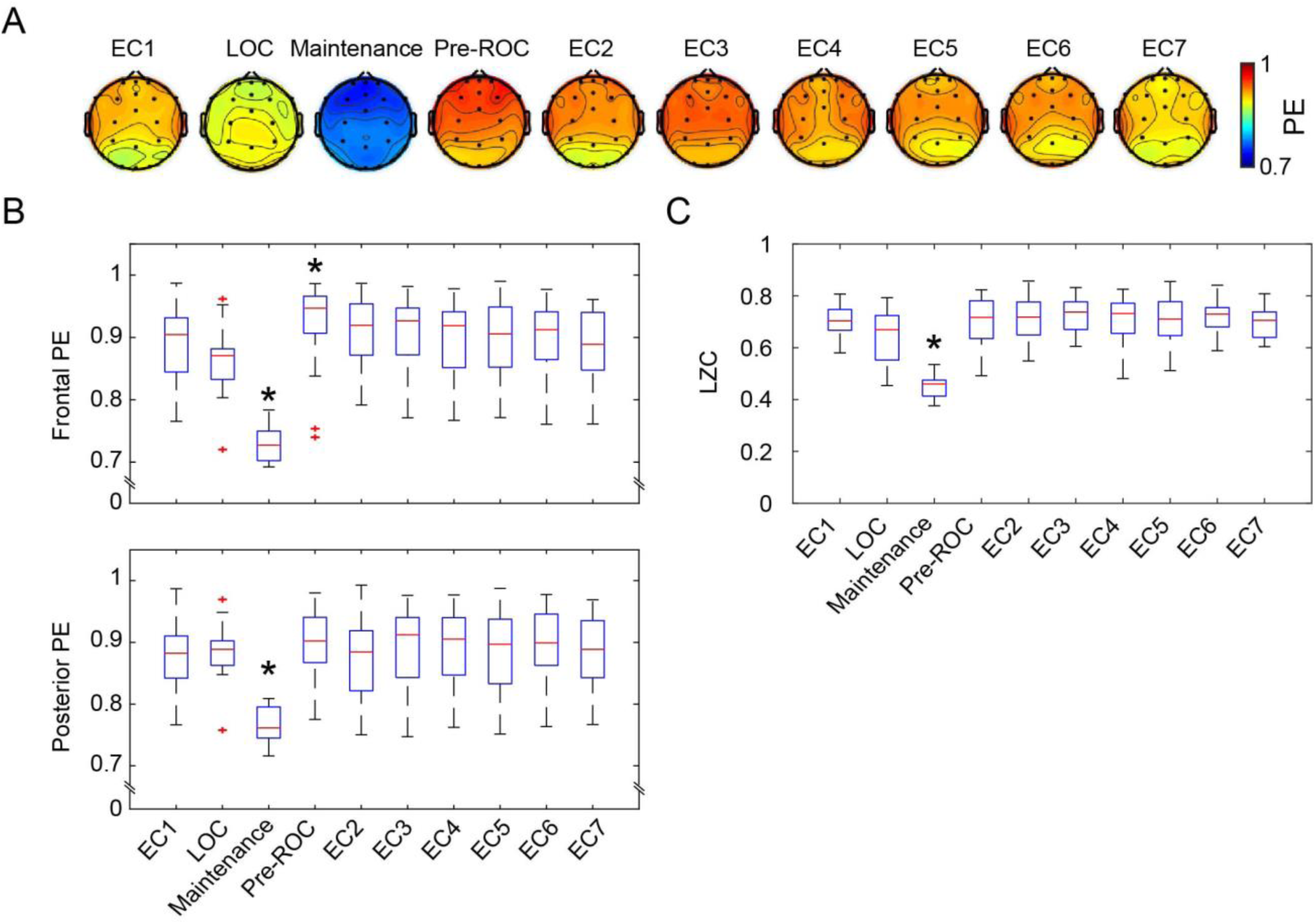
Cortical dynamics before, during, and after general anesthesia. (A) Scalp topographic maps of the group-level permutation entropy (PE; median average across N=30 participants) at the ten studied epochs. (B) The box plots of average PE values in frontal (Fp1, Fp2, Fpz, F3, F4 and Fz) and posterior channels (P3, P4, Pz, O1, O2 and Oz) for the studied epochs. On each box, the central mark is the median, the edges are the 25th and 75th percentiles, the whiskers extend to the most extreme data points determined by the MATLAB algorithm to be non-outliers, and the points deemed by the algorithm to be outliers are plotted individually (red cross). (C) The box plots of LZC values for the studied epochs. EC=eyes-closed resting state (EC1 is baseline consciousness, EC2-7 are post-emergence just prior to cognitive testing), LOC=loss of consciousness, Pre-ROC=2-minute epoch just before recovery of consciousness. *indicates significant difference relative to EC1, using linear mixed model analysis (Bonferroni-corrected p <0.05).

### Sleep-wake activity in the days following exposure to general anesthesia

Average rest-activity patterns for all study subjects are shown in **Figure 5A**. As expected, time of day significantly affected inactivity in both groups (F_129, 6867_ = 46.24, p < 0.0001). Cosinar analysis demonstrated that peak inactivity occurred between 3 and 4 am for both groups. Importantly, two-way ANOVA analysis revealed that over the week prior to the study day, there were no significant differences between the participants who would subsequently be anesthetized and those who would not (F_1, 6865_ = 3.70, p > 0.05). Moreover, there were no significant interactions between time and group (F_149, 6865_ = 1.18, p > 0.05). Analysis of rest-activity resumed upon completion of the experiment on the study day. Actigraphy revealed a small yet statistically significant effect of anesthetic exposure, accounting for only 0.19% of the total variance (F_1, 3576_ = 12.55, p = 0.0004). This was attributable to an increase in inactivity in the isoflurane-exposed volunteers during the initial two hours of the early evening (**Fig. 5B**); activity patterns were not distinguishable after that time period. As before the study day, time remained as a highly significant predictor of inactivity accounting for 42% of the variance (F_77, 3576_ = 35.78, p < 0.0001).

**Figure 5:**
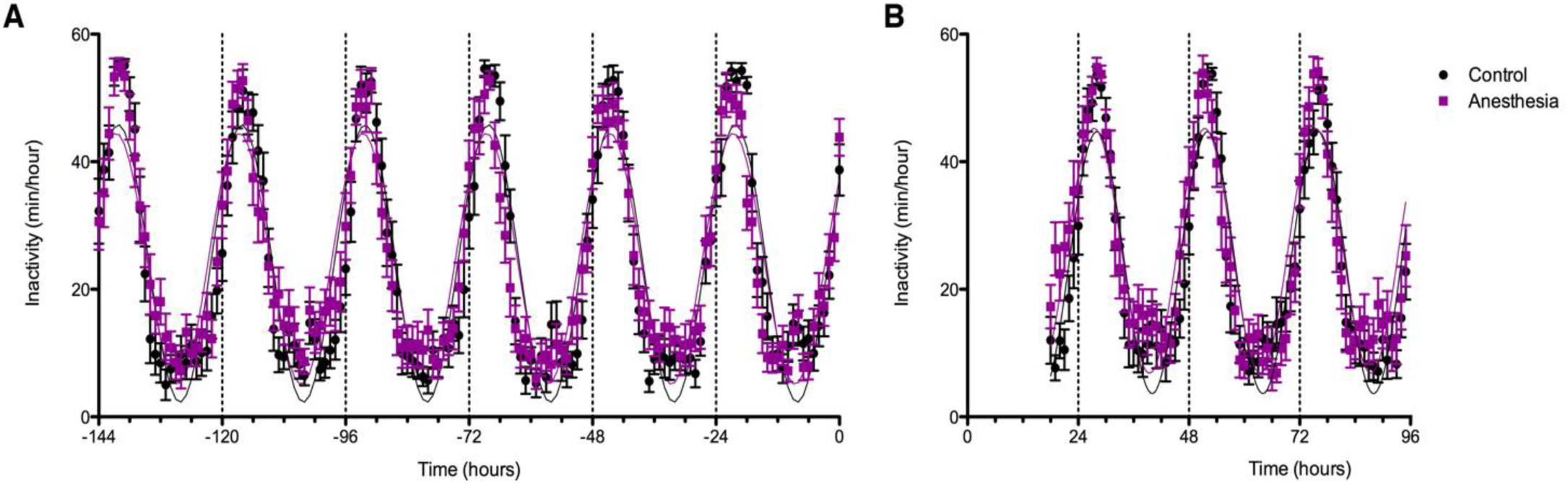
Effects of anesthetic exposure on rest-activity rhythms. (A) Rest activity plots are displayed in the week prior to the study day for volunteers that were subsequently randomised to anesthetized (purple) or control (black) conditions. (B) Rest-activity rhythms in the same participants are displayed on the evening of the study day and for the ensuing days. Time = 0 corresponds to midnight on the evening of the study day.

## Discussion

This the most comprehensive and best-controlled study assessing cognitive recovery from the anesthetized state in healthy humans. We have demonstrated that reconstitution of cognition after general anesthesia is a process that unfolds over time rather than a discrete event occurring all at once. Cognitive recovery also follows a counterintuitive sequence. Executive function was found to recover early, whereas tests of processing speed, attention, and reaction time had a more prolonged recovery. Furthermore, EEG-based measures of cortical dynamics return to baseline just prior to the recovery of consciousness after general anesthesia, with permutation entropy in frontal cortex statistically significantly higher than posterior cortex just after emergence. Finally, sleep-wake activity patterns are essentially unperturbed after anesthetic exposure.

This study is consistent with our prior analysis of source-localized alpha oscillations and graph-theoretical variables such as network efficiency and modularity, which were identified in a subset of participants who had high-density EEG and which also recovered within three hours after emergence from general anesthesia (Blain-Moraes et al., 2017). From a clinical perspective, it is remarkable that within three hours of recovery after a prolonged and deep general anesthetic, participants were performing a variety of complex cognitive tasks with similar accuracy and speed in comparison to participants who had not been anesthetized. On a shorter timescale, both local and global dynamic measures of cortical activity returned to baseline levels just prior to the return of responsiveness after general anesthesia. On a longer timescale, there was no evidence of disrupted sleep in the days following anesthetic exposure compared to non-anesthetized controls. This latter finding is striking considering that actigraphy measures of sleep are sensitive to stress, low alcohol exposure, and even sedative effects of non-alcoholic beer (Mezick et al., 2009; Franco et al., 2012; Geoghegan et al., 2012). The data suggest that, in healthy humans, networks supporting both higher cognition and arousal states recover uneventfully after deep general anesthesia.

The recovery of cortical dynamics in frontal-parietal networks just prior to the return of consciousness in our experimental paradigm should be considered in light of prior studies focused on emergence from general anesthesia. One study of healthy volunteers anesthetized with either propofol or dexmedetomidine and imaged using positron emission tomography found that those responsive to command exhibited activation of subcortical arousal centers with only limited frontal-parietal involvement (Langsjo et al., 2012). This is consistent with a more recent study of functional magnetic resonance imaging demonstrating a transient activation of brainstem loci upon emergence from propofol sedation (Nir et al., 2019) as well as a positron emission tomography study revealing subcortical and posterior cortical activation in association with the reversal of propofol sedation using the acetylcholinesterase inhibitor physostigmine (Xie et al., 2011). However, it is important to note that all of these studies involved experimental conditions in which there was ongoing exposure to a sedative-hypnotic, coupled with pharmacological or behavioral stimulation. By contrast, our paradigm reflects spontaneous emergence with residual isoflurane levels predicted to be 1 to 4 orders of magnitude below those required for hypnosis, which likely accounts for evidence of the robust return of cortical dynamics. However, it is worth noting that—in addition to the subcortical sites identified in human and animal studies (Kelz et al., 2019) —the prefrontal cortex might play a critical role in the control of arousal states of relevance to general anesthesia. A recent animal study demonstrated that cholinergic stimulation of medial prefrontal cortex, but not two sites in posterior parietal cortex, was sufficient to reverse the anesthetized state despite continuous administration of clinically relevant concentrations of sevoflurane (Pal et al., 2018). Thus, the prefrontal cortex might play a critical role in recovery of both consciousness (medial prefrontal) and cognition (dorsolateral prefrontal). Although causal inferences cannot be made based on the current study, the findings of early recovery of an executive function task known to involve dorsolateral prefrontal cortex (Berman et al., 1995) and a stronger recovery of frontal cortical dynamics after emergence from general anesthesia warrants further investigation.

Strengths of this study include: (1) surgically relevant anesthetic concentrations and duration of anesthetic exposure, in the absence of the confound of surgery itself; (2) multicentre design with substantially more participants than are typically found in studies of anesthetic-induced unconsciousness; (3) a non-anesthetized population to control for learning effects of cognitive assessment as well as comparison of sleep-wake patterns; and (4) complement of neurophysiological, cognitive, and behavioural measures on different time scales. Limitations of this study that constrain interpretation include: (1) young healthy population that precludes extrapolation of findings to older, younger, or sicker surgical patients encountered in clinical care; (2) only one anesthetic regimen tested, albeit a clinically common one, with unclear relevance to other perturbations such as sleep or pathological disorders of consciousness; (3) inability to blind participants to anesthetized and non-anesthetized conditions, which could potentially influence results; (4) relatively sparse EEG channels that were found to be a reliable sources of data across all participants; (5) no assessment of sleep macroarchitecture (i.e., rapid eye movement sleep vs. slow-wave sleep); (6) no longer-term cognitive or behavioural assessment beyond the first three days of post-anesthetic actigraphy; and (7) inability to assess source-localized brain regions, subcortical regions, or resting-state networks.

In conclusion, this study establishes neurophysiologic, cognitive, and behavioural recovery patterns after a surgically relevant general anesthetic in human volunteers. The rapid recovery of cortical dynamics just prior to recovering consciousness, the restored accuracy of executive function and multiple cognitive functions within three hours of emergence, and the normal sleep-wake patterns in the days following the experiment provide compelling evidence that the healthy brain is resilient to the effects of even deep general anesthesia. The findings also suggest that the immediate and persistent cognitive dysfunction identified after general anesthesia in healthy animals (Culley et al., 2004; Valentim et al., 2008; Carr et al., 2011; Callaway et al., 2012; Zurek et al., 2012; Jevtovic-Todorovic et al., 2013; Zurek et al., 2014; Jiang et al., 2017) does not necessarily translate to healthy humans and that postoperative neurocognitive disorders might relate to factors other than general anesthesia, such as surgery or patient comorbidity (Krause et al., 2019; Wildes et al., 2019).

## Materials and Methods

The full methods of this study (clinicaltrials.gov NCT01911195) have been published and are freely available (Maier et al., 2017).

### Ethics

This multicentre study was reviewed and approved by the Institutional Review Board specializing in human subjects research at the University of Michigan, Ann Arbor; University of Pennsylvania; and Washington University in St. Louis. Volunteers were recruited through the use of fliers and were compensated for their participation at levels approved by ethics committees. Participation eligibility required that all subjects provide written informed consent, which was obtained after careful discussion, in accordance with the Declaration of Helsinki.

### Experimental Design

The experimental design and data acquisition are summarized in **Figure 1**. This was a within-group study of anesthetized participants with a primary outcome of the pattern of cognitive recovery after general anesthesia; a non-anesthetized cohort was included to control for the learning effects of repeated cognitive testing and circadian factors. Volunteers were randomly assigned to receive general anesthesia with isoflurane or to engage in waking activity on the study day and serve as experimental controls. Baseline cognitive assessment was performed after an initial screening.

For anesthesia sessions, participants were closely monitored by two attending anesthesiologists during the study day. Attending anesthesiologists elicited a standard clinical preoperative history and physical examination, independently verified that volunteers met inclusion and fasting criteria, and safely conducted the anesthetic. Each subject underwent intravenous catheter placement. An appropriately fitted EEG head cap (Electrical Geodesics, Inc. Eugene OR) was affixed to the scalp. Electrical impedances on each channel were kept under 50kOhms/channel whenever possible. Standard anesthesia monitors (electrocardiogram, non-invasive blood pressure cuff, a pulse oximeter) were applied, and capnography was measured. Subjects completed a second baseline round of neurocognitive testing with ongoing EEG recordings. Upon completing the neurocognitive battery, subjects were pre-oxygenated by face mask prior to induction of general anesthesia with a stepwise increasing infusion rate of propofol: 100mcg/kg/min x 5 min, 200mcg/kg/min x 5 min, and then 300mcg/kg/min x 5 min. During this time, an audio loop issued commands every 30 seconds asking subjects to squeeze their left or right hand (in random order) twice. Loss of consciousness (LOC) was defined as the first time that a subject failed to respond to two sets of consecutive commands. After 15 minutes of propofol administration, subjects began inhaling isoflurane at 1.3 age-adjusted MAC (minimum alveolar concentration) (Nickalls and Mapleson, 2003). Thereafter, a laryngeal mask was inserted orally, a nasopharyngeal temperature probe was placed, and the propofol infusion was discontinued. Anesthetized subjects continued to inhale 1.3 age-adjusted MAC isoflurane anesthesia for three hours. Burst suppression, a sign of deep anesthesia, was found to be associated with this concentration of isoflurane in this cohort (Hemmings et al., 2019; Shortal et al., 2019). Blood pressure was targeted to remain within 20% of baseline pre-induction values using a phenylephrine infusion or intermittent boluses of ephedrine, as necessary. Pressure support ventilation was initiated with pressures titrated to maintain tidal volumes in the 5-8ml/kg range while end tidal carbon dioxide levels were targeted to 35-45 torr. Surface warming blankets were utilized to maintain body temperature in the normal range. Subjects received 4 mg intravenous ondansetron 30 minutes prior to discontinuation of isoflurane for antiemetic prophylaxis.

Isoflurane was discontinued at the end of the three-hour anesthetic period. Verbal command loops were reissued every 30 seconds upon completion of the isoflurane exposure. The laryngeal mask was removed when deemed medically safe by the attending anesthesiologists. Recovery of consciousness (ROC) was defined as the earliest instance in which subjects correctly responded to two consecutive sets of audio loop commands. At this point, defined as time = 0 minutes, subjects restarted neurocognitive testing with a brief pause between consecutive rounds. Neurocognitive testing was repeated at t= 30, 60, 90, 120, 150, and 180 minutes following emergence. Each battery of neurocognitive testing lasted approximately 15-25 minutes and was preceded by five minutes of eyes closed, resting state EEG data acquisition. Brief restroom or nutrition breaks were permitted between testing rounds, as necessary.

Subjects were discharged according to standard post-anesthesia care unit discharge criteria after completing their final battery of neurocognitive testing. A study site coordinator contacted each subject within 24 hours of the study day to document any adverse events.

A second group of healthy individuals (n=30) was recruited to participate in the same study design. These individuals also fasted overnight, but did not have intravenous lines inserted. Rather than being anesthetized, these volunteers remained awake (by reading or watching television on a personal electronic device) and continued fasting for 3.5 hours in order to control both for potential learning effects of repeated testing and also for circadian variability in testing performance (McLeod et al., 1982; Gur et al., 2001; Van Dongen et al., 2003; Van Dongen and Dinges, 2005; Jasper et al., 2009; Tucker et al., 2010). We chose not to sedate these participants as a control for the anesthetized state because doing so would have obscured the predicted learning effect accompanying repeated neurobehavioral testing and would thus confound the normal performance standard that was required. Volunteers randomised to the restful group were instructed to avoid napping and were regularly monitored by a dedicated research assistant.

### Participants

A total of 60, healthy American Society of Anesthesiologists physical status classification I or II volunteers were enrolled in the “Reconstructing Consciousness and Cognition” study (NCT01911195). The choice of study subject numbers was informed by several factors, including previous studies (Eger et al., 1997; Eger et al., 1998), biological plausibility (our best estimates regarding effect size and standard deviation), and safety considerations (exposing the minimum number of humans to general anesthesia in order to answer the questions of interest). The main factor that was considered in estimating our required sample size was the time difference in return of cognitive functions within the subjects receiving general anesthesia. Sample size calculation was modeled with various assumptions regarding the difference in recovery times between the first and last cognitive domains to return, and the standard deviations of these parameters. A range between 30 min and 90 min was considered for differences in recovery times between cognitive domains (possible effect sizes). A range between 20 min and 40 min was considered for standard deviations of these parameters. Assuming relatively conservative estimates (difference in recovery times = 30 min and standard deviation = 40 min), 30 subjects would provide >80% power with a two-sided alpha<0.05, using an unpaired t test.

With relatively liberal assumptions (difference in recovery times = 90 min and standard deviation = 20 min), 30 subjects would provide >99% power with a two-sided alpha<0.001, using an unpaired t test.

Each study site (University of Michigan, University of Pennsylvania, Washington University in St. Louis) recruited twenty volunteers who met the inclusion criteria. Prospective volunteers were screened using a phone questionnaire administered by a study coordinator. Eligible subjects that consented to participate in this study underwent a baseline familiarization round of neurocognitive testing and were given rest-activity monitoring devices (actiwatch) one week prior to the study day.

### EEG acquisition and analysis

To assess the neural correlates of the anesthetized state and recovery, participants enrolled at the University of Pennsylvania and University of Washington in St. Louis (n=40) were fitted with 32 EEG scalp electrodes. Subjects at the University of Michigan (n=20) were fitted with 64 or 128 EEG scalp electrodes. EEG recordings began prior to the baseline neurocognitive testing on the study day and were continued with minimal interruption until the completion of the final neurocognitive test.

#### EEG analysis

The raw EEG signals were exported into MATLAB (version 2015a; MathWorks, Inc., Natick, MA), down-sampled to 250 Hz (resample.m function in Matlab signal processing toolbox), and re-referenced to the linked-mastoid reference. Electrodes on the lowest parts of the face and head were removed, leaving 21 channels on the scalp (common to EEG montage for all participants) for the analysis. Data segments with obvious noise or non-physiological artefacts were identified and removed by visual inspection of the waveform and spectrogram of the EEG signals. Prior to the analysis, the EEG signals were bandpass filtered at 0.5 - 30 Hz via butterworth filter of order 4 (butter.m and filtfilt.m in MATLAB signal processing toolbox) to remove the possible baseline drift and muscle artefacts.

#### Analysis Epochs

Ten 2-min epochs were selected during the 7 resting-state eyes-closed sessions, and also during the exposure to anesthesia: 1) LOC - the first 2 minutes after loss of responsiveness; 2) Maintenance – the last 2 minutes before the discontinuation of isoflurane; and 3) Pre-ROC – the 2 minutes immediately preceding recovery of responsiveness. Burst-suppression patterns were present in the Maintenance epoch for 6 participants; to prevent the confounding effect of the suppression pattern on the EEG measures, we instead extracted 2-min continuous, non-suppression epochs in the last 10 minutes before the discontinuation of isoflurane (n=3 participants), or 7 minutes immediately after the discontinuation of isoflurane (n=3 participants), which showed similar spectral properties when compared to the other participants. Detailed information on the EEG sample size is listed in Table S1. For completeness, seven 2-min epochs were selected during the 7 resting-state eyes-closed sessions in the non-anesthetized group.

### Permutation Entropy

We used permutation entropy (PE) to measure the local dynamical changes of EEG in frontal and posterior channels. PE quantifies the regularity structure of a time series, based on a comparison of the order of neighbouring signal values, which is conceptually simple, computationally efficient, and artefact resistant (Bandt and Pompe, 2002), and has been successfully applied to the separation of wakefulness from unconsciousness (Li et al., 2008; Olofsen et al., 2008; Jordan et al., 2013; Ranft et al., 2016). The calculation of PE requires two parameters: embedding dimension (*d*_*E*_) and time delay (τ). In line with previous studies, we used *d*_*E*_=5 and τ=4 in order to provide a sufficient deployment of the trajectories within the state space of the EEG beta activity during wakefulness and anesthesia (Jordan et al., 2013; Ranft et al., 2016). Supplementary analysis was performed to test the sensitivity of PE using alternative strategies of parameter selection.

In the implementation, each 2-min epoch was divided into non-overlapping 10-sec windows, the PE was calculated for each window, and the PE values were averaged across all the windows for each studied epoch and channel. The topographic maps of group-level PE value for each studied epoch was constructed using the topoplot function in the EEGLAB toolbox (Delorme and Makeig, 2004). For statistical comparisons, the averaged PE values were calculated over the frontal (Fp1, Fp2, Fpz, F3, F4 and Fz) and posterior channels (P3, P4, Pz, O1, O2 and Oz) at each studied epoch for each participant.

#### Lempel-Ziv Complexity

Lempel-Ziv Complexity (LZC) was computed as a surrogate of complexity to reflect the spatiotemporal repertoire across scalp potentials. LZC is a method of symbolic sequence analysis that measures the complexity of finite length sequences (Lempel and Ziv, 1976), which has been shown to be a valuable tool to investigate brain states (Casali et al., 2013; Abasolo et al., 2015; Schartner et al., 2015; Hudetz et al., 2016; Schartner et al., 2017). The calculation of LZC requires a binarization of the multichannel EEG data. In this study, we used the implementation as described in (Schartner et al., 2015; Schartner et al., 2017), and calculated the instantaneous amplitude from the Hilbert transformed EEG signal for each channel, which was binarized using its mean value as the threshold for the current channel (supplementary analysis was performed to test the effect of threshold selection). The data segment was then converted into a binary matrix, in which rows represent channels and columns represent time points, capturing the complexity or diversity in both temporal and spatial domains. LZC was computed by rearranging the binary matrix time point by time point, searching the resultant sequence for the occurrence of consecutive characters, or “words” and counting the number of times a new word is encountered (Hudetz et al., 2016).

For implementation, the average signal was first subtracted from all channels in order to remove the effect of common reference, and then the multichannel EEG epochs were divided into non-overlapping 4-sec windows to compute the LZC, with the resultant LZC values being averaged across all the windows for each studied epoch. In line with previous studies, we normalized the original LZC by the mean of the LZC values from N=50 surrogate data generated by randomly shuffling each row of the binary matrix, which is maximal for a binary sequence of fixed length (Schartner et al., 2015; Schartner et al., 2017) (supplementary analysis was performed to test alternative methods in the generation of surrogate data).

#### Statistical analysis of EEG measures

Statistical analyses were conducted in consultation with the Center for Statistical Consultation and Research at the University of Michigan. All EEG-derived PE and LZC values were exported to IBM SPSS Statistics version 24.0 for Windows (IBM Corp. Armonk, NY). Statistical comparisons were performed using linear mixed models (LMM), to test (1) the difference between the ten studied epochs for both PE and LZC measures, and (2) the difference between PE values derived from frontal and posterior channels. In contrast to traditional repeated-measures ANOVA analysis, LMM analysis offers more flexibility in dealing with missing values (see Table S1) and accounting for the within-participant variability by including a random intercept associated with each participant. The non-anesthetized control group was included primarily to control for learning effects in repeated cognitive testing and thus, for EEG analysis, the statistical analysis was focused on anesthetized group. For the model of LZC values, the fixed effect is the studied epoch. For the model of PE values, the fixed effects include the studied epoch, region, and the interaction between them. We fitted the models with random intercept specific for each participant and used the default variance components as the covariance structure. We modelled the studied epoch as repeated effects and assumed each studied epoch was associated with different residual variance by using the diagonal structure as the covariance structure of the residuals. We employed restricted maximum likelihood estimation. The models described above were chosen by taking into account the information criteria and likelihood ratio test results in the comparisons, with alternative models including additional random effects and repeated effects, as well as different covariance structures (Table S2). For all post hoc pairwise comparisons, the Bonferroni corrected p-value along with the estimate and 95% confidence interval (CI) of the difference were reported. A two-sided p < 0.05 was considered statistically significant.

### Neurocognitive testing

Neurocognitive tests were selected from the Cognition test battery (Basner et al., 2015) to reflect a broad range of cognitive domains, ranging from basic abilities such as sensory-motor speed to complex executive functions such as abstraction. The order of the six tests was randomised but balanced across subjects. Individual subjects took the tests in the same order (except during familiarization). In each test session, subjects repeated the first test after completion of sixth test. Therefore, the temporal resolution for one test in five control subjects and five experimental subjects was doubled. The following six tests, adopted from (Basner et al., 2015), were chosen for this study.

The *Motor Praxis Task (MP)* measures sensorimotor speed and validates that volunteers have sufficient command of the computer interface. Participants were instructed to click on squares that appear randomly on the screen, with each successive square smaller and thus more difficult to track. The test depends upon function of visual and sensorimotor cortices (Gur et al., 2001; Gur et al., 2010; Neves et al., 2014).

The *Psychomotor Vigilance Test (PVT)* measures a volunteer’s reaction times (RT) to visual stimuli that are presented at random inter-stimulus intervals over 3 minutes (Basner et al., 2011). Subjects monitor a box on the computer screen, and press the space bar once a millisecond counter appears and begins timing response latency. In the well-rested state, or whenever sustained attention performance is optimal, right frontoparietal cortical regions are active during this task. Conversely, with sleep deprivation and other suboptimal performance, studies demonstrate increased activation of default-mode networks during this task, which is considered to be a compensatory mechanism (Drummond et al., 2005).

The *Digit-Symbol Substitution Task (DSST)* adapts the Wechsler Adult Intelligence Scale (WAIS-III) for a computerized presentation. The DSST required participants to refer to a continuously displayed legend that matches each numeric digit to a specific symbol. Upon presentation of one of the nine symbols, subjects must select the corresponding number as rapidly as possible. The DSST primarily recruits the temporal cortex, prefrontal cortex, and motor cortex. Activation of frontoparietal cortices during DSST performance has been interpreted as reflecting both on-board processing in working memory and low-level visual search (Usui et al., 2009).

The *Fractal 2-Back (F2B)* is an extremely challenging nonverbal variant of the Letter 2-Back test. N-back tests probe working memory. The F2B consisted of the sequential presentation of a set of fractals, each potentially repeated multiple times. Participants were instructed to respond when the current stimulus matched the stimulus previously displayed two images ago. The F2B is well-validated task that robustly activates dorsolateral prefrontal cortex, cingulate, and hippocampus (Ragland et al., 2002).

The *Visual Object Learning Test (VOLT)* measures the volunteer’s memory for complex figures (Glahn et al., 1997). Participants memorize 10 sequentially displayed three-dimensional figures. Subsequently, they were instructed to select the familiar objects that they memorized from a larger set of 20 sequentially presented objects that included the 10 memorized and 10 similar but novel objects. Visual object learning tasks have been shown to depend upon frontal and bilateral anterior medial temporal cortices as well as the hippocampus(Jackson and Schacter, 2004).

#### Statistical analysis of cognitive data

Apart from the Bayesian analyses, statistical analyses were implemented by the SAS software version 9.4. In view of the learning effect with repeated cognitive testing, it is difficult to pinpoint when any cognitive domain returns to baseline. We could therefore not simply compare recovery times between individual cognitive tests, as we had planned. Instead, in order to test whether there was a difference in recovery times between cognitive domains, we opted for a Bayesian regression approach using all the data from the anesthetized subjects as well as the non-anesthetized controls. For this analysis, we used the brms package in R (R Foundation for Statistical Computing, Vienna, Austria), which uses Stan for full Bayesian inference. We simulated M samples from the (posterior) conditional distribution of the model parameters given the data, and for each set of simulated model parameters, we calculated the corresponding recovery time, thus we obtained M samples of recovery times. Then, based on these simulated recovery times, we further calculated the corresponding differences in recovery times between pairs of cognitive tests (or domains), and evaluated the posterior probability of one cognitive test recovering more than 30 minutes before another test [P(diff>30 minutes|data)] by checking the sample proportion in the posterior sample. Regarding priors, we chose normal priors with a large variance, which is a routine choice for non-informative flat priors.

Multiple statistical comparisons were separately conducted on the standardized speed and accuracy indices, which were two metrics to evaluate each task performance. We used nonlinear mixed-effects models (NLMM) based on a damped exponential in time to fit the data of each task at all time-points, i.e. *y*_*t*_ = *y*_*baseline*_ *+ α + β* ∗ *e*^*γt*^, where *y*_*t*_ is the task performance response at time *t, y*_*baseline*_ is the pre-treatment baseline response, *α* is the random intercept, *β* is the random slope, and *γ* is the coefficient of the fixed-effect time *t*. The random intercept *α* and the random slope *β* are independent, and distributed from normal distributions. Some NLMMs were degenerated to models with a fixed intercept or a fixed slope according to goodness of fit. The appeal of NLMM analysis is its flexibility in modelling the nonlinear trend of cognitive data with repeated measures over time, and the ability to adjust the pre-treatment baseline performance. In order to test against the alternative hypothesis that there was a significant difference at the end of 3-hour period between the anesthetized group and the controlled group, we performed model-based multiple testing on the least squares means of predicted response difference at 3 hours. For speed and accuracy of each task, the point estimates of the difference as well as the Bonferroni corrected confidence intervals (CI) of the difference with an overall significance level 0.05 were reported. The P-values of testing difference between groups should be compared to the corrected level 0.05/6=0.0083.

### Actigraphy

In order to assess and control for differences in baseline sleep-wake rhythms and to potentially evaluate the effect of isoflurane anesthesia on subsequent rest activity behaviour, participants were trained and instructed to wear a wrist GT3X+ device (ActiGraph) on their nondominant wrist beginning at the conclusion of their baseline visit, one week before prior to and one week following their assigned study day. Actigraphy data were downloaded to a computer and GT3X+ devices recharged on the study day and again at completion one week following the study day. Raw activity counts for each subject were binned into 1-minute epochs and analysed for bouts of inactivity using the Cole-Kripke scoring algorithm (Cole et al., 1992), included in the ActiLife 6.7.2 software. For each subject, minutes of inactivity each hour were calculated. ActiLife’s wear time validation was employed using default settings to confirm that subjects used the GT3X+ monitor as instructed. Hours in which wear time validation revealed that the watch was not worn, were excluded from the analysis.

#### Statistical analysis of actigraphy data

Actigraphy data were imported into Prism 5.0d (GraphPad) and analysed with a two-way ANOVA with Time in hours relative to the midnight before testing and Treatment Group (Isoflurane exposed or Awake Controls) as the two factors. Effects of Time, Treatment group, and the interaction between these two factors were considered to be significant for P values < 0.05. Due to asynchrony in times during which the GT3X+ device was not worn across individuals, it was not possible to conduct a repeated measures two-way ANOVA. To obtain a graphical best fit of actigraphy data, we performed a standard Cosinar analysis.

## Acknowledgments

This study was funded by a collaborative grant from the James S. McDonnell Foundation, St. Louis, MO; National Institutes of Health (Bethesda, MD, USA) grant T32GM112596; and the anesthesiology departments of the University of Michigan, University of Pennsylvania, and Washington University.

