## Supplementary Figures-Table for "Recovery of Consciousness and Cognition after General Anesthesia in Humans"

**Supplementary Figure 1.** Confirmative results of permutation entropy (PE) with different settings of parameters of embedding dimension ($d_E$) and time delay ($\tau$). The description of the figure and statistical results are same with those in Figure 4 A-B.

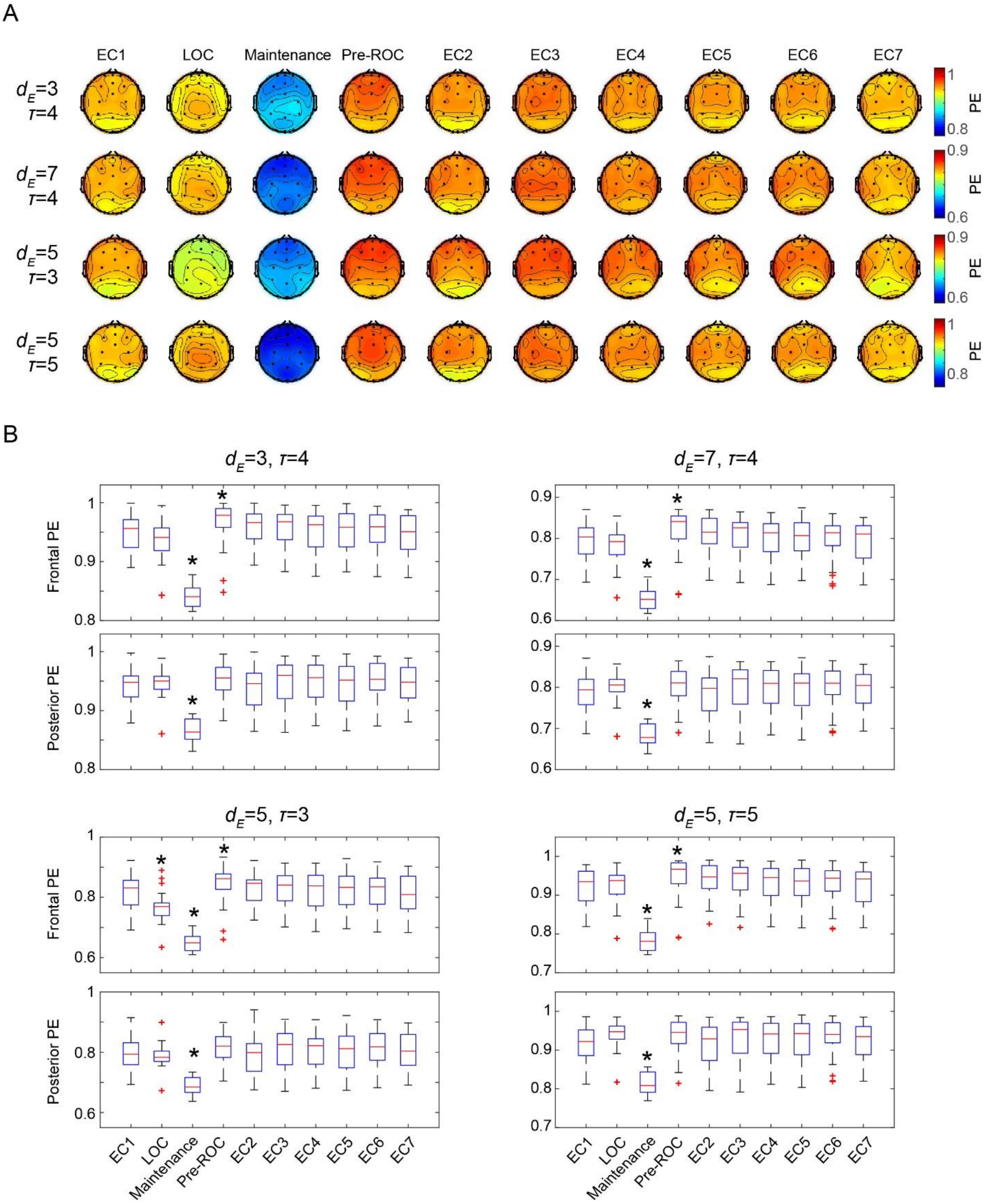

**Supplementary Figure 2.** Confirmative results of Lempel-ziv Complexity (LZC). (A) Sensitivity of the LZC measure to the selection of threshold for binalization. (B) Confirmative results of LZC by using spatial shuffling (top) and phase randomization (bottom) in the generation of surrogate data for normalization. The spatial shuffling method permutates the spatial order (at each time point) of the spatiotemporal matrix. The phase randomization method preserves the spectral profile of the signals and reflects the complexity changes beyond the power spectrogram. The description of the figure and the statistical results are same with those in Figure 4 C.

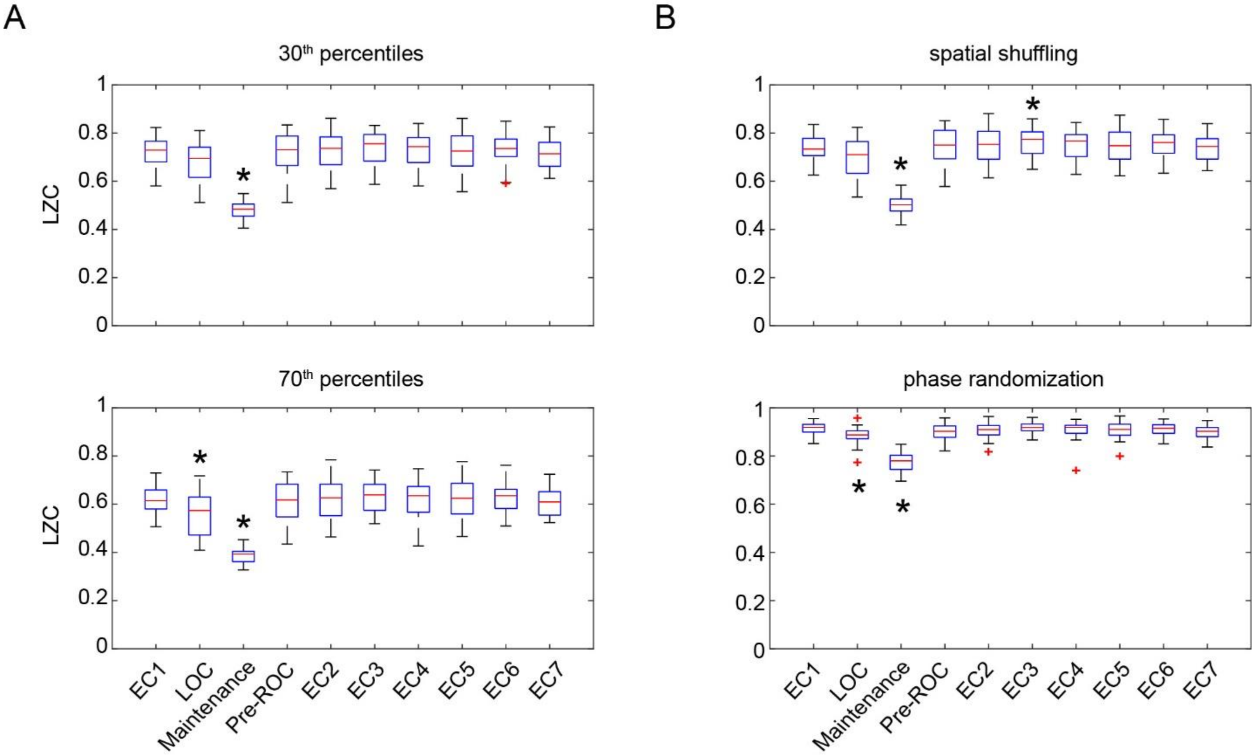

**Supplementary Figure 3.** Cortical dynamics as assessed by PE and LZC for the non-anesthetized control group. (A) Scalp topographic maps of the group-level PE at the seven resting-state eyes-closed (EC) epochs. (B) The box plots of average PE values in frontal and posterior channels for the studied epochs. (C) The box plots of LZC values for the studied epochs.

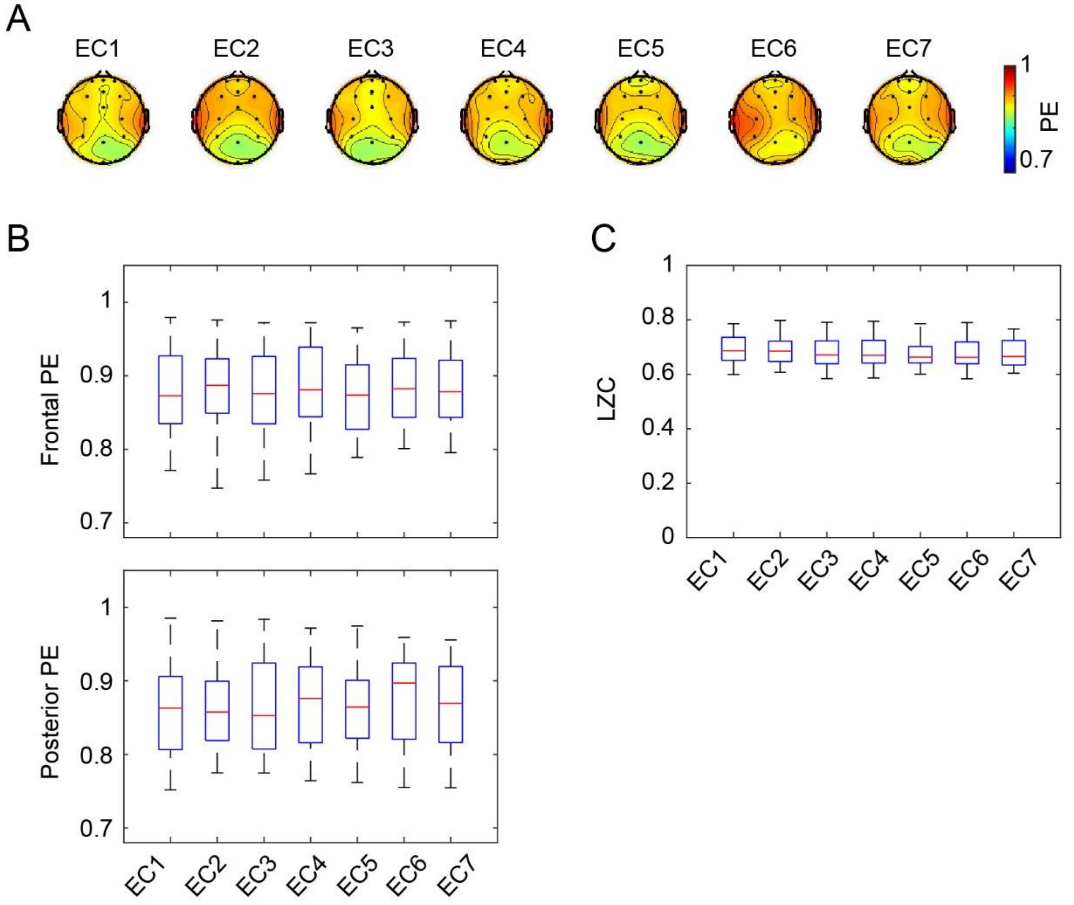

**Supplementary Table 1.** Sample size of data epochs in EEG analysis

| Epochs | Anesthetized group | Control group |
|---|---|---|
| EC1 | n=29 | n=28 |
| LOC | n=30 | -- |
| Maintenance | n=29 | -- |
| Pre-ROC | n=28 | -- |
| EC2 | n=28 | n=30 |
| EC3 | n=28 | n=30 |
| EC4 | n=29 | n=29 |
| EC5 | n=29 | n=30 |
| EC6 | N=30 | n=30 |
| EC7 | N=29 | n=30 |

Sample size indicates the number of participants whose data were available for each studied epoch. For EC sessions, the missing data are because the sessions were not recorded; for the anesthetic epochs, the EEG recordings were temporarily interrupted during maintenance (n=1) or during emergence (n=1).

**Supplementary Table 2.** Model selection for the statistical analysis of EEG measures

| EEG measures | LMM models | Number of parameters | AIC | BIC | -2 Restricted Log Likelihood | LST statistics |
|---|---|---|---|---|---|---|
| PE | The employed model | 31 | -2143.831 | -2096.263 | -2165.831 | -- |
| | The model with random center effect | 33 | -2145.203 | -2088.986 | -2171.203 | $\chi^2(2)$=5.372, p=0.068 |
| | The model with region as additional repeated effect | 41 | -2133.480 | -2042.669 | -2175.480 | $\chi^2(10)$=9.649, p=0.472 |
| | The model with AR1 as the covariance structure of the residuals | 23 | -2073.166 | -2060.193 | -2079.166 | $\chi^2(8)$=86.665, p<0.001 |
| LZC | The employed model | 21 | -779.126 | -739.182 | -801.126 | -- |
| | The model with random center effect | 23 | -779.732 | -732.526 | -805.732 | $\chi^2(2)$=4.606, p=0.100 |
| | The model with AR1 as the covariance structure of the residuals | 13 | -714.628 | -703.734 | -720.628 | $\chi^2(8)$=80.498, p<0.001 |

LMM, linear mixed model; AIC, Akaike's Information Criterion; BIC, Schwarz's Bayesian Criterion; LST, Likelihood Ratio Tests; AR1, first-order autoregressive

The information criteria (AIC, BIC) are displayed in smaller-is-better form, i.e. a smaller value of the criterion indicates a “better” fit. The LSTs are based on comparing the values of likelihood functions (-2 Restricted Log Likelihood) between the employed model and the model with varied covariance structures of random-effects and residuals, which under mild regularity conditions asymptotically follows a $\chi^2$ distribution, with the number of degrees of freedom as obtained by the difference in the number of parameters of the two models.

The model with random center effect assumes that the subjects in each experimental site/center are associated with different variance. Both PE and LZC data seem obtain a better fit with the employed model without center-specific effect, as reflected by the non-significant LST statistics (this test model vs. the employed model) and the smaller values of BIC.

The model of PE that considers the region as an additional repeated effect resulted in larger values in AIC and BIC, and the non-significant LST statistic (this test model vs. the employed model) implied a better fit with the employed model without repeated region effect.

The model with AR1 as the covariance structure of the residuals uses a smaller number of parameters for both PE and LZC data, but the smaller values in AIC and BIC, and the significant LST statistic (the employed model vs. this test model) suggested a better fit with the employed model that used diagonal covariance structure for the residuals.
